# Intrinsically photosensitive retinal ganglion cell mediated pupil function is impaired in Parkinson’s disease

**DOI:** 10.1101/169946

**Authors:** Daniel S. Joyce, Beatrix Feigl, Graham Kerr, Luisa Roeder, Andrew J. Zele

## Abstract

Parkinson’s disease is characterised by non-motor symptoms including sleep and circadian disruption, but the underlying aetiology is not well understood. Melanopsin-expressing intrinsically photosensitive Retinal Ganglion Cells (ipRGC) transmit light signals from the eye to brain areas controlling circadian rhythms and the pupil light reflex. Here we evaluate the hypothesis that these non-motor symptoms in people with Parkinson’s disease may be linked to ipRGC dysfunction. Using chromatic pupillometry, we measured intrinsic (melanopsin-mediated) ipRGC and extrinsic (rod/cone photoreceptor-mediated) inputs to the pupil control pathway in a group of optimally medicated participants with a diagnosis of Parkinson’s disease (PD, *n* = 17) compared to controls (*n* = 12). Autonomic tone was evaluated by measuring pupillary unrest in darkness. The PD participants underwent additional clinical assessments using the Unified Parkinson’s disease Rating Scale (UPDRS) and the Hoehn and Yahr scale (H&Y).

Compared to controls, the PD group demonstrated an attenuated pupil constriction amplitude in response to long wavelength pulsed stimulation, and reduced post-illumination pupil response (PIPR) amplitude in response to both short wavelength pulsed and sinusoidal stimulation. In the PD group, PIPR amplitude did not correlate with measures of sleep quality, retinal nerve fibre layer thickness, UPDRS or H&Y score, or medication dosage. Both groups exhibited similar pupillary unrest in darkness.

We show that melanopsin and the rod/cone-photoreceptor contributions to the pupil control pathway are impaired in people with early-stage Parkinson’s disease. Given that the deficits are independent of clinical assessment severity and are observed despite optimal medication, the melanopsin-mediated PIPR may be a biomarker for the detection of Parkinson’s disease and its continued monitoring in both medicated and unmedicated individuals.

## Introduction

Parkinson’s disease (PD) is a debilitating disorder characterised by a loss of dopamine (DA) producing neurons in regions of the basal ganglia, impairing autonomic function and resulting in motor symptoms including tremor, rigidity, and bradykinesia (Chaudhuri *et al*., Jankovic, 2008). By the time these symptoms manifest, up to 60% of dopaminergic cells within the substantia nigra pars compacta are destroyed (Dauer and Przedborski, 2003). Non-motor symptoms can precede motor symptoms and include sleep disturbances and daytime sleepiness, fatigue, depressed mood and cognitive impairments (Chaudhuri *et al.*, Pagan, 2012). Due to their earlier onset, these symptoms may have clinical utility as early biomarkers of the disease (Chaudhuri *et al.*, Pagan, 2012).

The aetiology underlying sleep and circadian disturbances in Parkinson’s disease is not well understood, but is hypothesised to include dysregulation of the circadian system due to reduced dopaminergic neurotransmission (for review see Videnovic and Golombek, 2013). In people with Parkinson’s disease, a 4-fold reduction in melatonin expression has been observed without altered circadian phase (Videnovic *et al*., 2014), while in mouse models of the disease suprachiasmatic nucleus (SCN) signalling is reduced. These studies suggest degradation of environmental light signal processing via the retinohypothalamic tract that projects from the retina to the SCN.

In humans, the origin of the retinohypothalamic tract is a novel class of photoreceptors in the eye called intrinsically photosensitive retinal ganglion cells (ipRGCs) (Dacey *et al*., 2005, Liao *et al*., 2016, Nasir-Ahmad *et al*., 2017). IpRGCs make up less than 0.5% of all retinal ganglion cells (Liao *et al*., 2016, Nasir-Ahmad *et al*., 2017) yet project to over a dozen brain areas including those involved in circadian photoentrainment, sleep and mood regulation, and the pupil light reflex (Provencio *et al*., 1998, Gooley *et al*., 2001, Berson *et al*., 2002, Hattar *et al*., 2002, Dacey *et al*., 2005, Hattar *et al*., 2006, Baver *et al*., 2008, Do *et al*., 2009, Hannibal *et al*., 2014). The transmission of light signals to the brain by ipRGCs is initiated at two sites within the retina, either intrinsically via the endogenous melanopsin photopigment expressed in the ipRGC body (soma and dendrites) in the inner retina (Hattar *et al*., 2002, Provencio *et al*., 2002, Belenky *et al*., 2003, Do *et al*., 2009) or via extrinsic (synaptic) input from rod and/or cone photoreceptors in the outer retina (Dacey *et al*., 2005) that also involve dopaminergic amacrine intermediary cells (Belenky *et al*., 2003, Zhang *et al*., 2008, Van Hook *et al*., 2012, Hu *et al*., 2013). Melanopsin is maximally light sensitive in the short wavelength (blue) region of the visible spectrum and its physiological response is characterised by slow temporal kinetics and sustained signalling after light cessation (Dacey *et al*., 2005); in humans, the kinetics of the pupil light response after stimulus offset (the Post-Illumination Pupil Response, PIPR) provide a signature, non-invasive measure of melanopsin function (Gamlin *et al*., 2007, Markwell *et al*., 2010, Adhikari *et al*., 2015, Adhikari *et al*., 2016) that can be differentiated from extrinsic photoreceptor inputs using non-invasive chromatic pupillometry (see *Stimuli* and *Experimental Paradigms and Analyses*) (Kardon *et al*., 2009, Park *et al*., 2011, Feigl and Zele, 2014). Pupillometric assessment of ipRGCs in humans has shown clinical promise for a range of retinal and non-retinal diseases (for review see Feigl and Zele (2014)); pupil constriction in response to light stimuli has been used to evaluate outer retinal rod/cone dysfunction in Parkinson’s disease (Stergiou *et al*., 2009) but intrinsic ipRGC-mediated pupil function has not been investigated in Parkinson’s disease. Our primary aim was to determine if ipRGC function is impaired in people with Parkinson’s disease by performing chromatic pupillometry on optimally medicated Parkinson’s disease participants compared to a healthy age-matched control group.

It is well established that autonomic nervous system function is impaired in Parkinson’s disease (Dewey, Visser *et al*., 2004), and the resting pupil diameter is set by the autonomic nervous system through a dynamic equilibrium between parasympathetic input to the pupillary sphincter and sympathetic input to the dilator muscle (Lowenstein *et al*., 1963, McDougal and Gamlin, 2015). Early research in unmedicated Parkinson’s patients showed increased pupil diameters after light adaptation, reduced pupil constriction amplitude and prolonged time to pupil constriction (Micieli *et al*., 1991). In a group of mostly (71%, *n* = 12) unmedicated Parkinson’s disease patients the pupillary unrest in darkness was increased (Jain *et al*., 2011). To evaluate the level of autonomic tone in optimally mediated people with Parkinson’s disease, the secondary aim was to measure pupillary unrest in the absence of light stimulation.

## Materials and Methods

### Participants

Twenty-nine participants were recruited, comprising of 17 people with Parkinson’s disease (mean age = 64.9 years, SD = 6.1) and 12 control participants (mean age = 59.7 years, SD = 4.1). As shown in Table 1, participants with Parkinson’s disease were assessed as early stage with a mild to moderate disease severity (Unified Parkinson’s Disease Rating Scale (Fahn *et al*., 1987, Ramaker *et al*., 2002); Hoehn & Yahr (Hoehn and Yahr, 1967)) and were independent and cognitively intact (Mini-Mental State Examination (Folstein *et al*., 1975). All people with Parkinson’s disease were optimally medicated during all measurements (Table 1).

**Table 1.** Parkinson’s disease characterisation.

|  | Gender | Age<br>(years) | LEDD | MMSE | ACE-R | UPDRS | H&Y |
| --- | --- | --- | --- | --- | --- | --- | --- |
| Participant |  |  |  |  |  |  |  |
| PD1 | F | 59 | 998 | 30 | 98 | 34 | 1.5 |
| PD2 | M | 63 | 750 | 30 | 89 | 20 | 1 |
| PD3 | M | 66 | 400 | 30 | 87 | 45 | 2.5 |
| PD4 | M | 71 | 1064 | 29 | 89 | 30 | 2 |
| PD5 | M | 56 | 400 | 29 | 90 | 16 | 2 |
| PD6 | M | 74 | 1222.5 | 30 | 97 | 38 | 2.5 |
| PD7 | F | 69 | 400 | 29 | 98 | 24 | 2 |
| PD8 | F | 65 | 933 | 29 | 91 | 32 | 2 |
| PD9 | M | 66 | 225 | 30 | 89 | 26 | 2 |
| PD10 | M | 72 | 300 | 28 | 79 | 44 | 2 |
| PD11 | M | 63 | 525 | 26 | 75 | 58 | 2.5 |
| PD12 | M | 64 | 475 | 30 | 94 | 33 | 1 |
| PD13 | F | 56 | 612.5 | 29 | 98 | 35 | 1 |
| PD14 | M | 57 | 400 | 29 | 98 | 34 | 1 |
| PD15 | M | 70 | 0 | 28 | 89 | 43 | 2 |
| PD16 | F | 59 | 450 | 30 | 100 | 60 | 1 |
| PD17 | M | 74 | 400 | 29 | 91 | 45 | 1 |
| Mean |  | 64.9 | 597.2 | 29.1 | 91.3 | 36.3 | 1.7 |
| SD |  | 6.1 | 302.1 | 1.1 | 6.9 | 12.0 | 0.6 |

Note: LEDD = Levodopa equivalent daily dosage in mg; MMSE = Mini-mental state examination; ACE-R = Addenbrooke's cognitive examination - revised; UPDRS = Unified Parkinson's disease rating scale; H&Y = Hoehn and Yahr scale. Participant PD15 was not medicated at the time of participation and is excluded from the mean and SD calculation.

A comprehensive ophthalmic examination was completed in all Parkinson’s disease and control participants. All participants had a best corrected visual acuity ≥ 6/6 (Bailey-Lovie Log MAR Chart) and no ocular pathology on slit lamp examination or ophthalmoscopy. Intraocular pressure measured with non-applanation tonometry (iCare, Finland Oy, Helsinki, Finland) was within the normal range (< 21 mmHg) before dilation and at the conclusion of testing. All participants had normal trichromatic colour vision as assessed by the Farnsworth D-15. Participants with intra-ocular lenses were excluded from participation in this study and all participants had clear lenses.

Retinal nerve fibre layer thickness was measured using Optical Coherence Tomography (OCT) (Cirrus-HD OCT, Carl Zeiss Meditec, Inc., Dublin, CA, USA and Nidek RS-3000 RetinaScan Advance, Nidek Co., Ltd., Tokyo, Japan). Given the evidence for sleep disturbances in people with Parkinson’s disease and that ipRGCs mediate the environmental light signals for circadian photoentrainment, sleep quality was assessed in all participants using the Pittsburgh Sleep Quality Index questionnaire (PSQI) (Buysse *et al*., 1989).

All experimental protocols were approved by the University Human Research Ethics Committee and participants provided informed consent in accordance with the tenets of the Declaration of Helsinki.

### Pupillometer

Light stimuli were generated using a custom built extended Maxwellian-view optical system (Beer *et al*., 2005, Kankipati *et al*., 2010, Joyce *et al*., 2015, Joyce *et al*., 2016). The light from two 5 mm diameter LEDs (short wavelength, ‘blue’ appearing light, λ_max_ = 465 nm; full width half maximum (FWHM) = 19 nm; long wavelength, ‘red’ appearing light, λ_max_ = 638 nm, FWHM = 15 nm) was imaged in the plane of the pupil via two Fresnel lenses (100 mm diameter, 127 mm and 70 mm focal lengths; Edmund Optics, Singapore) and a 5° light shaping diffuser (Physical Optics Corp., California USA) which generated a 35.6° stimulus light. The consensual pupil response was recorded with a Pixelink camera (IEEE-1394, PL-B741 FireWire; 640x480 pixels; 60 frames.s^−1^) through a telecentric lens (Computar 2/3″ 55 mm and 2× Extender C-Mount) under infrared LED illumination (λ_max_ = 851 nm). A chin rest, temple bars and a head restraint maintained alignment in Maxwellian-view. Custom software coded in Matlab (version 7.12.0, Mathworks, Massachusetts USA) controlled stimulus presentation, pupil recording and analysis. Details of the pupillometry measurements are given elsewhere (Feigl *et al*., 2011, Zele *et al*., 2011).

### Stimuli

Short wavelength stimuli included predominant intrinsic ipRGC inputs to the pupil control pathway shown to be mediated by the melanopsin photopigment (Gamlin *et al*., 2007, Markwell *et al*., 2010, Adhikari *et al*., 2015). Long wavelength stimuli served as a measure of activity biased to the extrinsic outer retina photoreceptor activity and thus as a control stimulus that has minimal intrinsic ipRGC (melanopsin-mediated) activation. The light stimulation protocol consisted of a 10 s pre-stimulus baseline recording, pulsed (8 s rectangular) or phasic (12 s, 0.5 Hz sinusoidal) stimulus presentation, and a 40 s post-stimulus recording period (see Fig. 1A,B & C,D for the stimulus protocol). Short and long wavelength stimulus irradiances were equated to 15.1 log photons.cm^-2^.s^-1^ at the cornea.

**Figure 1.**
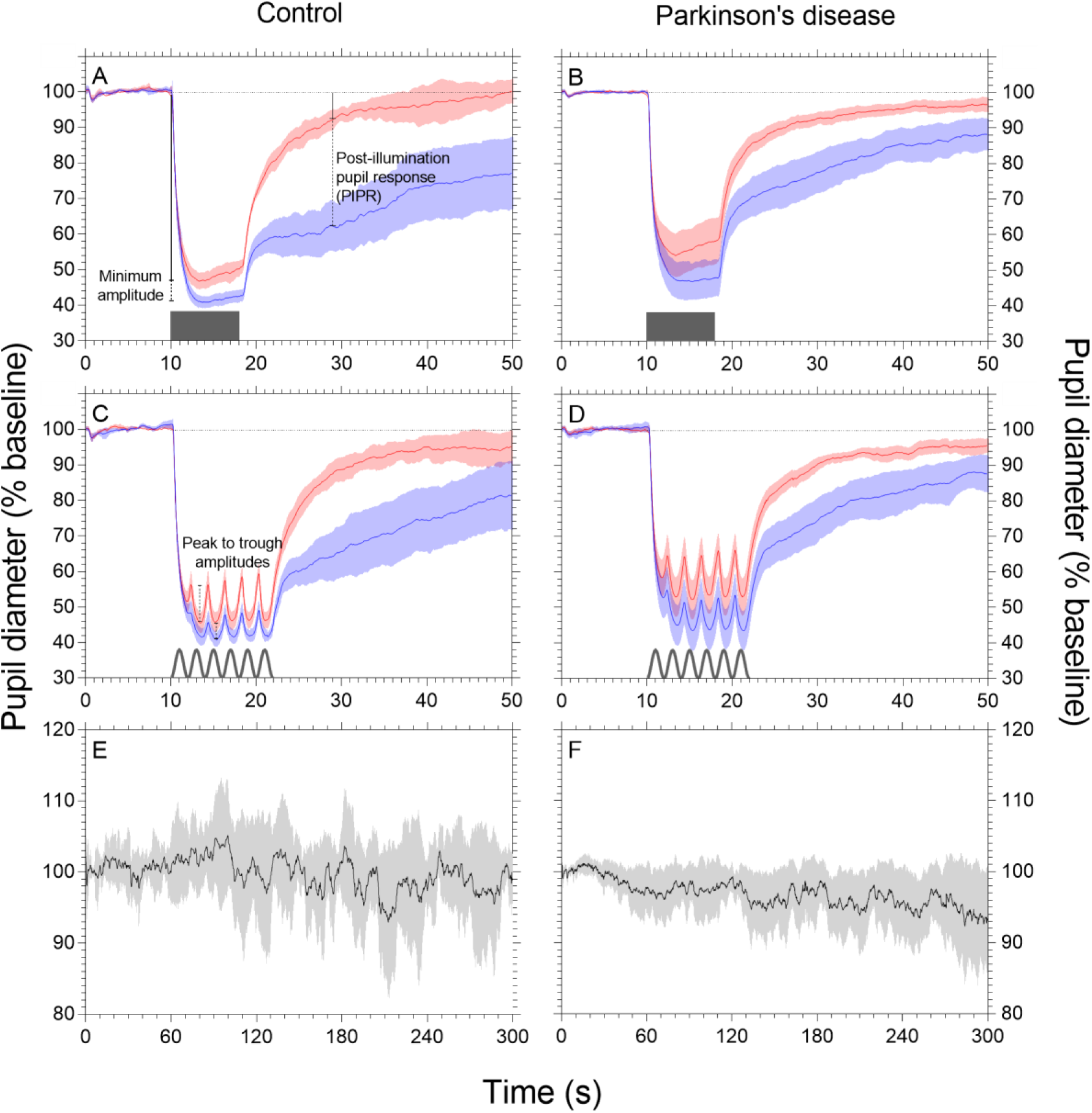
Normalised mean pupil waveforms during pulsed and sinusoidal stimulation, and pupillary unrest. Left panels show the control data (*n* = 12), right panel show the data for the participants with Parkinson’s disease (*n* = 17). The pupil metrics are illustrated in Panels A and C (minimum constriction amplitude, PIPR amplitude and peak to trough amplitude). A schematic of the test stimuli are depicted on the abscissa in the upper panels (pulsed stimuli) and middle panels (sinusoidal stimuli). The mean unrest data are shown in Panels E and F. Blue, red and grey shadings indicate 95% confidence intervals. To control for individual differences in baseline pupil diameter, the data are normalised to the first 10 s of recording.

Given the older age of the participants, retinal irradiances were estimated based upon established corrections for age-related changes in the optical density of the media of the eye (cornea, lens, aqueous and vitreous humours) for stimuli greater than 3° in diameter (van de Kraats and van Norren, 2007). It was calculated that the average attenuation by the optical media of the short wavelength stimuli was 0.54 log units in the PD group and 0.50 log units in the control group. The optical media attenuated the long wavelength stimuli by 0.16 log units for both groups. Long wavelength stimulation was therefore a control condition invariant to group membership and age and autonomic reactivity. To account for the proposed bistability of melanopsin (Mure *et al*., 2009) and participant fatigue (*Kankipati et al., 2010*, Feigl *et al*., 2011) stimuli were alternated, beginning with the long wavelength stimulus followed by the short wavelength stimulus. Two recordings for each wavelength of the pulsed and sinusoidal were obtained and the data report the average response for each condition. Pupillary unrest (see *Experimental Paradigms and Analyses* section) was recorded in the dark for 5 minutes at the end of the pulsed and sinusoidal testing to measure autonomic tone and fatigue. Each participant therefore underwent a total of 9 trials during a recording period lasting approximately 1.5 hours.

### Experimental Paradigms

In order to investigate the interaction between inner and outer retina photoreceptor inputs to the pupil control pathway during pulsed stimulation, constriction amplitude was measured (Kardon *et al*., 2009, Joyce *et al*., 2015). To determine the interaction between inner and outer retinal contributions to the phasic pupil response of the dark adapted pupil, two parameters were estimated – the peak to trough amplitude (Joyce *et al*., 2015), and the Phase Amplitude Percentage [PAP: (long wavelength peak to trough – short wavelength peak to trough) / long wavelength peak to trough] (Feigl and Zele, 2014). To assess intrinsic melanopsin signalling, the post-illumination pupil response amplitude can be measured at any time >1.7 s after stimulus offset (Adhikari *et al*., 2016). The melanopsin-mediated PIPR under short wavelength conditions demonstrates a sustained constriction (that is, a reduction from baseline diameter) that is the signature of melanopsin activity signalled via the intrinsic ipRGC pathway. In contrast, the PIPR amplitude to long wavelength stimulation is less sustained and rapidly returns to baseline due to the low sensitivity of melanopsin to long wavelength light (Dacey *et al*., 2005, Kardon *et al*., 2009). We calculated the optimal timing of the PIPR metric given our equipment, sample, and stimulus conditions: Using the control group data only, the pulsed and sinusoidal PIPR data were averaged within the short and long wavelength conditions. Subtracting the short from long wavelength data determined the timing of the largest difference between these retinal inputs to the PIPR, which was the 1 s window (Park *et al*., 2011) of the 11th second after light offset. Thus the PIPR value used for all analyses (both PD and control groups) was 11 s after light offset.

Parkinson’s disease is characterised by changes in autonomic tone (Goetz *et al*., 1986, Micieli *et al*., 2003), whereby the balance of sympathetic and parasympathetic systems is impaired. Because the dilator and sphincter pupil muscles that maintain the steady-state pupil diameter receive sympathetic and parasympathetic innervation respectively (for review see (McDougal and Gamlin, 2015)), we measured pupil diameter in the dark for 5 minutes in order to quantify changes in the spontaneous oscillations of the pupil (i.e. pupil unrest) during this period, which may differ with disease status. We used Fourier analysis to calculate metrics of RMS, dominant frequency (Hz), dominant frequency (dB), and approximate entropy (Pincus, 1991, Morrison *et al*., 2008). Data were analysed in Matlab 2016a (The MathWorks, Inc., Natick, Massachusetts, USA). We also calculated the average pupillary unrest index (PUI; (Lüdtke *et al*., 1998)) for each individual, over a shortened duration of five minutes in order to minimise fatigue because it was conducted at the end of the experimental session. The PUI is an additive measure of consecutive pupil diameters to quantify pupil oscillation instability (Lüdtke *et al*., 1998).

### Statistical Analysis

Each pupil tracing was individually visualised and blinks were linearly interpolated in Matlab. In order to minimise the correlations between the pupil light reflex metrics when expressed in millimetres (Joyce *et al*., 2016), the data were normalised to the average pupil diameter of the first 10 seconds and expressed as percentage baseline units. The non-normally distributed data for the PD and control groups were compared using independent samples Mann-Whitney U tests. Correlations within the PD group data were explored using Spearman’s rank order test. All statistical analyses were performed in SPSS Statistics (v23.0, IBM, Armonk, NY, USA) using two-tailed tests with an alpha level of *p*<.05. Participant data are reported using box plots that demonstrate the *median*, *interquartile range*, *maximum and minimum*.

### Procedure

Participants with Parkinson’s disease were assessed for disease severity (UPDRS, H&Y) and cognitive impairment (MMSE) prior to visual testing. All participants were provided the PSQI and instructed in its use (sent via mail and returned on the day of testing), to assess their quality of sleep in the four weeks prior to visual testing. Upon presentation participants had a comprehensive ophthalmic exam, before dilation of their stimulated eye (Tropicamide 0.5% w/v, Bausch & Lomb). Once the pupil had fully dilated the participant was briefed of the protocols and aligned in the pupillometer. All pupillometry was conducted in the dark and before each trial participants adapted to the dim room illumination (< 1 lux) for 7 minutes. Between trials the participants were permitted to remove their head from the pupillometer but remained seated. Following pupillometry, participants had their fundus and lens examined (slit lamp), retinal nerve fibre layer thickness measured via OCT, and IOP re-assessed. The entire experimental and ophthalmic testing was completed within two hours.

## Results

The Optical Coherence Tomography measurements of the optic disc retinal nerve fibre layer thickness were similar between the PD group (*median* = 93.00 μm, *IQR* = 19.50) and control group (89.50 μm, 21.00) (*p* = .902). Sleep quality as measured by the PSQI was reduced in the PD group (7.00, 4.00) compared to controls (4.00, 3.00), but this difference was not significant (*p* = .264) and groups did not differ along derived 2-factor dimensions of sleep quality (*p* = .517) and sleep efficiency (*p* = .578) (Magee *et al*., 2008).

The pupil light reflex (*mean ±95% confidence intervals*) for the control (left panels) and PD (right panels) groups in response to the pulsed and sinusoidal stimuli are shown in Fig. 1A-D). Fig. 1A,B,C,D demonstrate the reduced PIPR amplitude during long compared to short wavelength stimulation. The pupillary unrest waveforms (*mean ±95% confidence intervals*) are shown for the control and PD group participants in Fig. 1E and F, respectively.

Box plots (Fig 2) show all participant data for the minimum amplitude (pulsed stimuli) and PIPR amplitude pupil metrics (pulsed and sinusoidal stimuli). The minimum pupil constriction amplitude for short wavelength pulsed stimulation was similar between the PD (*median* = 43.35%, *interquartile range* = 10.57%) and control groups (39.93%, 2.79%) (*p* = .079), whereas the minimum constriction amplitude for long wavelength pulsed stimulation was reduced in the PD group (51.48%, 9.73%) compared to the control group (46.10%, 6.05%) (*p* = .034). The melanopsin-mediated PIPR amplitude was measured in the pulsed and sinusoidal pupillometry protocols. For short wavelength stimuli that have a high melanopsin excitation, the pulsed PIPR amplitude was 14.73% higher in Parkinson’s disease participants (80.32%, 23.16%) compared to controls (65.59%, 20.52%) (*p* = .018), indicating reduced melanopsin contributions to this process (i.e., closer to baseline diameter in the PD group than controls). Similarly, short wavelength sinusoidal PIPR amplitude was 12.96% higher in the PD group (81.72%, 15.21%) compared to controls (68.76%, 21.32%)(*p* = .011). As expected, the long wavelength (with minimal melanopsin excitation) PIPR amplitude was not different between groups for either pulsed (*p* = .325) or sinusoidal (*p* = .556) stimulation.

**Figure 2.**
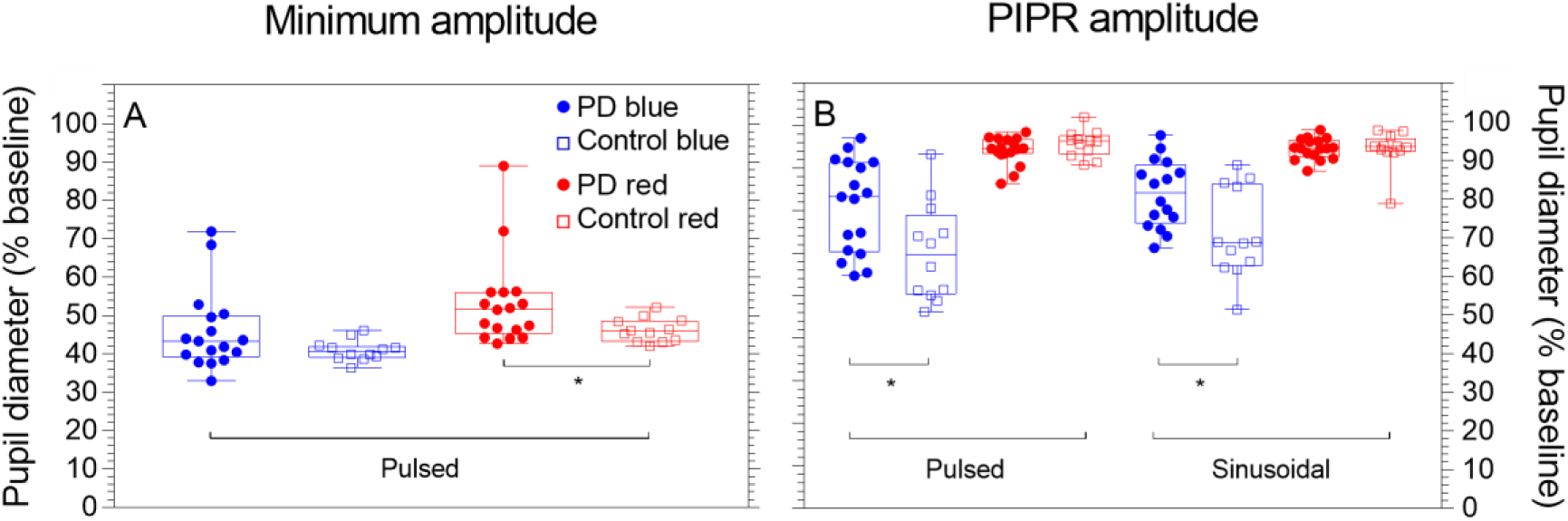
Minimum amplitude and post-illumination pupil response (PIPR) amplitude in response to pulsed and sinusoidal stimulation. Each data point represents an individual’s mean data, boxplots depict the quartiles and whiskers the range. Asterisks indicate a significant difference between groups (*p* < .05).

Spearman’s rank-order correlations were performed to determine if the short wavelength pulsed PIPR amplitude was associated in the PD group with sleep quality (PSQI), symptom severity (UPDRS), retinal nerve fibre layer thickness (RNFL) or medication dosage (LEDD); no statistically significant correlations were observed (Table 2).

**Table 2.** Spearman’s rank-order correlations between pulsed short wavelength PIPR amplitude and Parkinson’s disease markers.

|  | PIPR | RNFL | UPDRS | LEDD | PSQI |
| --- | --- | --- | --- | --- | --- |
| PIPR | 1 | .107 (.682) | .136 (.602) | .241 (.352) | .259 (.315) |
| RNFL | .107 (.682) | 1 | -.248 (.337) | -.172 (.509) | .072 (.783) |
| UPDRS | .136 (.602) | -.248 (.337) | 1 | -.113 (.666) | .118 (.651) |
| LEDD | .241 (.352) | -.172 (.509) | -.113 (.666) | 1 | .477 (.053) |
| PSQI | .259 (.315) | .072 (.783) | .118 (.651) | .477 (.053) | 1 |

Note: Data are expressed as *correlation coefficient (p value)*. PIPR = Post-illumination pupil response, RNFL = Retinal nerve fibre layer thickness, UPDRS = Unified Parkinson's Disease Rating Scale, LEDD = Levodopa equivalent daily dosage, PSQI = Pittsburgh sleep quality index. $n = 17$ .

In response to sinusoidal stimulation, the peak to trough amplitude and the phase amplitude percentage (PAP) of the phasic pupil response (Fig. 3) shows more variability in participants with Parkinson’s disease than controls, independent of stimulus wavelength. With short wavelength lights that have high melanopsin excitation (Fig. 3A) the peak to trough amplitude trended to increase in the PD group (7.95%, 3.57%), which is indicative of reduced melanopsin contributions compared to controls (5.59%, 2.20%), but this difference was not significant (*p* = .205). Similarly, under long wavelength stimulation with low melanopsin excitation (Fig. 3A), the peak to trough amplitude did not differ between the PD group (12.03%, 6.41%) and controls (11.48%, 3.18%) (*p* = .471). The median PAP did not significantly different between groups (*p* = .537; Fig. 3B).

**Figure 3.**
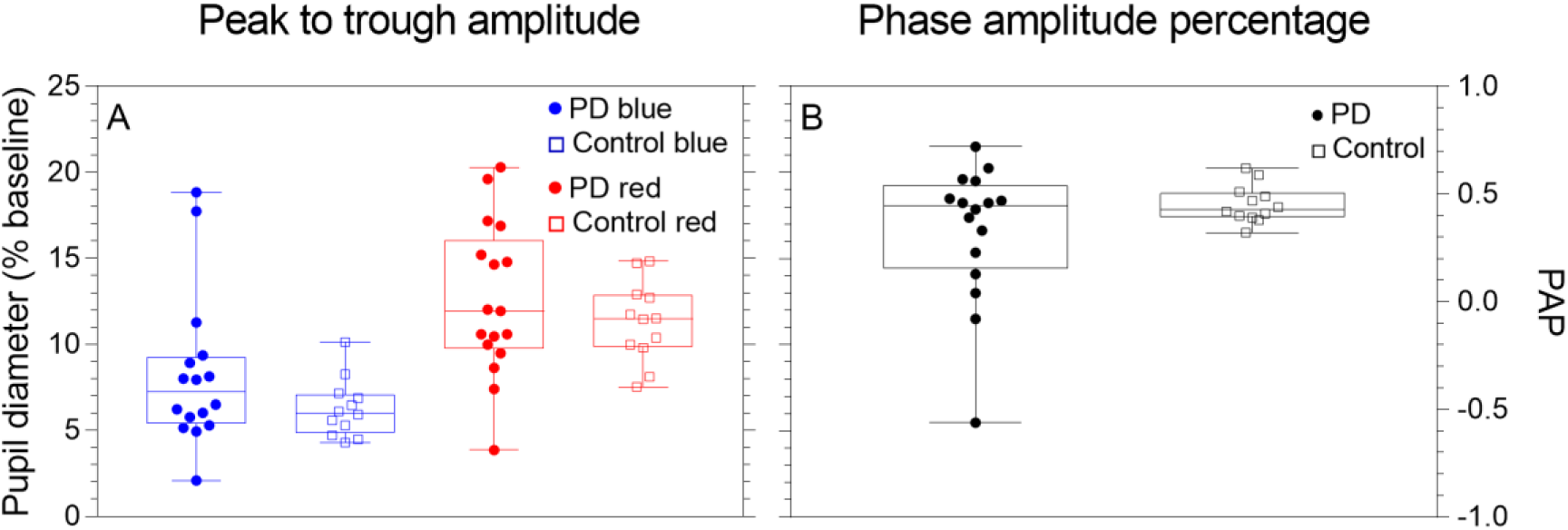
Peak to trough amplitude and phase amplitude percentage derived during sinusoidal stimulation.

Pupillary unrest (see Fig. 1E,F for mean normalised waveforms) measured autonomic tone and fatigue. The results of the Fourier analysis and pupillary unrest index (PUI) of each individual tracing are given in Table 3; the PD and control groups did not statistically differ on any metric.

**Table 3.**
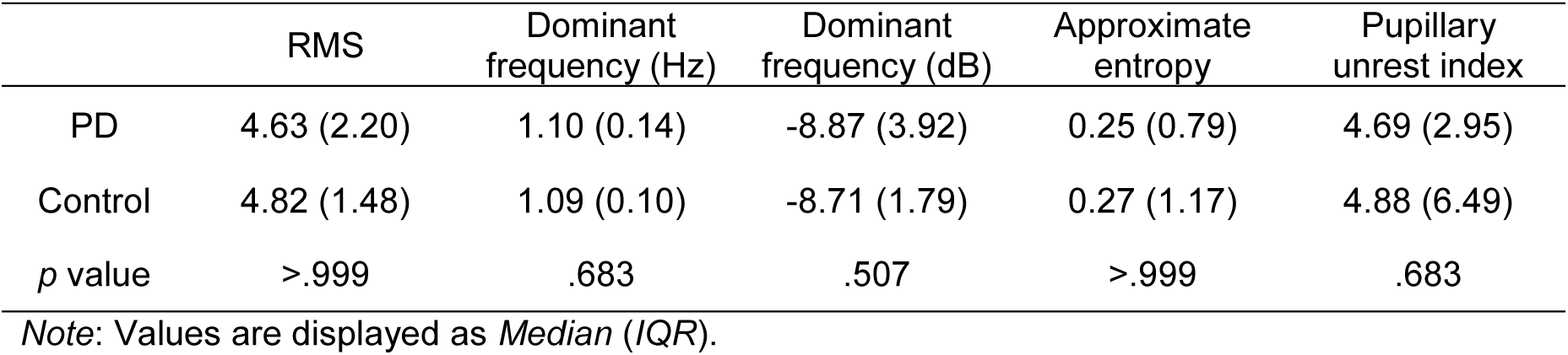
Medians and interquartile ranges of the pupillary unrest metrics.

## Discussion

This study investigated whether pupil control differed in optimally medicated individuals with Parkinson’s disease compared to healthy age-matched controls. Deficits in the PD group relative to controls were observed in the post-illumination pupil response to short wavelength stimulation and the constriction amplitude in response to long wavelength stimulation. Pupillary unrest was not significantly different between groups, neither was there a significant sleep deficit as assessed with the PSQI.

The reduction in the post-illumination pupil response amplitude (closer to baseline diameter) in the PD group compared to controls indicates that melanopsin-mediated ipRGC inputs to pupil control pathway are impaired, and that this effect size is both large and clinically relevant (difference between medians = 17.49%). Reduced ipRGC function has been associated with impaired sleep in retinal diseases (Gracitelli *et al*., 2015, Maynard *et al*., 2017) and while there was reduced sleep quality in patients with Parkinson’s disease compared to the control group, this difference was not statistically significant. We acknowledge however that alternative methods of sleep assessment such as polysomnography may be more sensitive than the PSQI in detecting sleep deficits.

The PIPR amplitude was reduced in response to both pulsed and sinusoidal stimulation in the PD group, and these deficits were observed in the Parkinson’s disease participants with no retinal thinning. Previous studies have identified reduced retinal nerve fibre layer thickness in people with Parkinson’s disease including at the early-to mid-stage (Inzelberg *et al*., 2004, Hajee *et al*., 2009). That the PD group did not statistically differ in RNFL thickness compared to controls is consistent with the early stage diagnosis based upon their clinical UPDRS and H&Y scores (Kerr *et al*., 2010). Because ipRGCs are relatively few in number (~0.1 to ~ 0.4% of all retinal ganglion cells in human retinae (Liao *et al*., 2016, Nasir-Ahmad *et al*., 2017), deficits in function may become apparent before a reduction in gross ganglion cell numbers can be detected by RNFL thickness.

Given the aetiology of Parkinson’s disease, deficits in ipRGC function could be linked to a reduction in dopamine expression. IpRGCs form retinal circuits with dopaminergic amacrine cells and may themselves be sensitive to DA through feedback loops (Viney *et al*., 2007, Vugler *et al*., 2007, Zhang *et al*., 2008, Allen *et al*., 2014). The PIPR amplitude is reduced in patients with type II diabetes without diabetic retinopathy (Feigl *et al*., 2011), which in rodent models features decreased retinal dopamine (Nishimura and Kuriyama, 1985, Aung *et al*., 2014). Post-mortem examination reveals that DA cell morphology is abnormal in the Parkinson’s disease retina, with reductions in both DA and DA’s synthesising enzyme tyrosine hydroxylase (Nguyen-Legros, 1988, Djamgoz *et al*., 1997), but retinal DA is reduced for unmedicated but not medicated patients with Parkinson’s disease (Harnois and Di Paolo, 1990). The observed deficits in PIPR amplitude could therefore reflect a mechanism other than dopaminergic dysfunction because the PD group were optimally medicated, and PIPR amplitude was not correlated with measured disease severity characteristics (Table 2). Alternate hypotheses include deficiencies in the cholinergic inputs to the pupil control system (Fotiou *et al*., 2009), compatible with cholinergic gait disturbances in Parkinson’s disease (Rochester *et al*., 2012, Bohnen *et al*., 2013); or reduced ipRGC signaling due to α-synuclein deposition within the inner plexiform and ganglion cell layers (Beach *et al*., 2014, Bodis-Wollner *et al*., 2014).

The constriction response to long wavelength square wave stimulation, unaffected by yellowing of the lens with ageing, represents contributions of the extrinsic (rod and predominantly cone, due to their long wavelength sensitivity) ipRGC pathway. This pathway was impaired in the PD group compared to controls with a small but statistically significant difference (5.38%). Consistent with this, Micieli *et al.* (1991) used a light adapted paradigm (1200 Lux for 10 minutes) in unmedicated people with Parkinson’s disease and found slower pupil constriction latency and timing as well as a larger 12.58% reduction in constriction amplitude. Pupillometric deficits in outer retinal-mediated responses may parallel visual performance deficits in the Parkinson’s disease fovea (where the rods are absent), including colour vision, contrast sensitivity, and electroretinography (for review see Bodis-Wollner (2013).

Pupillary unrest metrics did not differ between the PD and control groups, exhibiting both low entropy (suggesting signal regularity) and similar dominant frequencies between groups. In contrast to this, Jain *et al.* (2011) reported that compared to controls, their largely (71%) unmedicated PD group with similar disease severity to our sample (H&Y = 1.7 (0.6), UPDRS = 20.5 (9.6)) had increased pupillary unrest using a longer 11-minute protocol. Medication may potentially influence the pupil control pathway that sets baseline pupil size, obscuring deficits in pupillary unrest mediated by the autonomic system (which balances the sympathetic and parasympathetic equilibrium), but the light-dependent PIPR drive can still demonstrate deficits in optimally medicated populations.

This study is the first to assess melanopsin-mediated ipRGC function in people with Parkinson’s disease. We demonstrate that the post-illumination pupil response, a marker of melanopsin pathway function, is disrupted in optimally medicated individuals with the disorder. Melanopsin dysfunction may therefore be a biomarker of Parkinson’s disease in both medicated and unmedicated individuals, with the potential to detect prodromal Parkinson’s given that the PIPR amplitude is uncorrelated with both clinical ratings of the disease and medication dosage. Future studies with larger sample sizes could further optimise the waveform, timing, and irradiance of stimulation in assessing deficits in the PLR in people with Parkinson’s, including longitudinal study to test the hypothesis that PIPR amplitude should increase (i.e. increased ipRGC dysfunction) with increasing disease duration. Given that ipRGCs are the primary conduit for photic entrainment to the solar day (Berson *et al*., 2002), and they innervate brain centres involved in sleep/wake regulation (e.g. SCN, ventrolateral preoptic area) (Hattar *et al*., 2006) their reduced function may play an important role in the pathophysiology of sleep and circadian rhythms in Parkinson’s disease.

## Acknowledgments

We thank Dr Prema Sriram, Mr Damien Lonergan and Mr Christopher Rogers for assistance.

## Author contributions

DSJ, BF, GK and AJZ designed the research.

DSJ, BF, LR, GK and AJZ performed data collection.

DSJ, BF, GK and AJZ performed data analysis and interpretation.

DSJ, BF, GK and AJZ prepared the manuscript.

